# Human genetic variations reveal Chromosomal Instability aiding Variants (CIVa) in kinetochore-microtubule associated proteins

**DOI:** 10.1101/2022.01.22.477339

**Authors:** Asifa Islam, Janeth Catalina Manjarrez-González, Trupti Gore, Xinhong Song, Viji M. Draviam

## Abstract

The vast majority of Chromosomal Instability (CIN) promoting mutations remain unknown. We assess the prevalence of Chromosomal Instability aiding Variants (CIVa) by collating Loss-of-Function (LoF) variants predicted in 135 chromosome segregation genes from over 150,000 humans, including consanguineous individuals. Surprisingly, we observe heterozygous and homozygous CIVa in Astrin and SKA3 genes that encode evolutionarily conserved microtubule-associated proteins essential for chromosome segregation. By combining high-resolution microscopy and controlled protein expression, we show the naturally occurring Astrin variant, p.Q1012*, as potentially harmful because it fails to localise normally, delays anaphase onset, induces chromosome misalignment and promotes chromosome missegregation. We show that N-terminal frameshift variants in Astrin and SKA3 are likely to generate shorter isoforms that do not compromise chromosome segregation revealing resilient mechanisms to cope with harmful variants. This study provides a framework to predict and stratify naturally occurring CIVa, an important step towards precision medicine for CIN syndromes.

## Main

Chromosomal Instability (CIN) is a hallmark of several pathologies^1^. CIN can arise from errors in chromosome-microtubule attachment events, leading to the loss or gain of chromosomes, disorganisation of nuclear structure, transcriptional heterogeneity and replication and proteotoxic stress^2–4^. Chromosome-microtubule attachment is facilitated by a macromolecular protein structure, the kinetochore (reviewed in ^5^). Variations in kinetochore (KT) proteins are known to cause genetic disorders including primary microcephaly (MCPH, OMIM 604321, OMIM 616051^6^) and mosaic variegated aneuploidy (MVA, OMIM 257300^7^). Although genetic disorders are prevalent in communities where consanguineous marriages are common^8–10^, there has been no systematic study of kinetochore gene variants in these communities. By combining live-cell microscopy with LoF genetic variant analysis, we set out a framework to identify and stratify the cellular and functional impact of <u>C</u>hromosomal Instability aiding <u>Va</u>riants (CIVa) (Fig. 1A).

**Figure 1.**
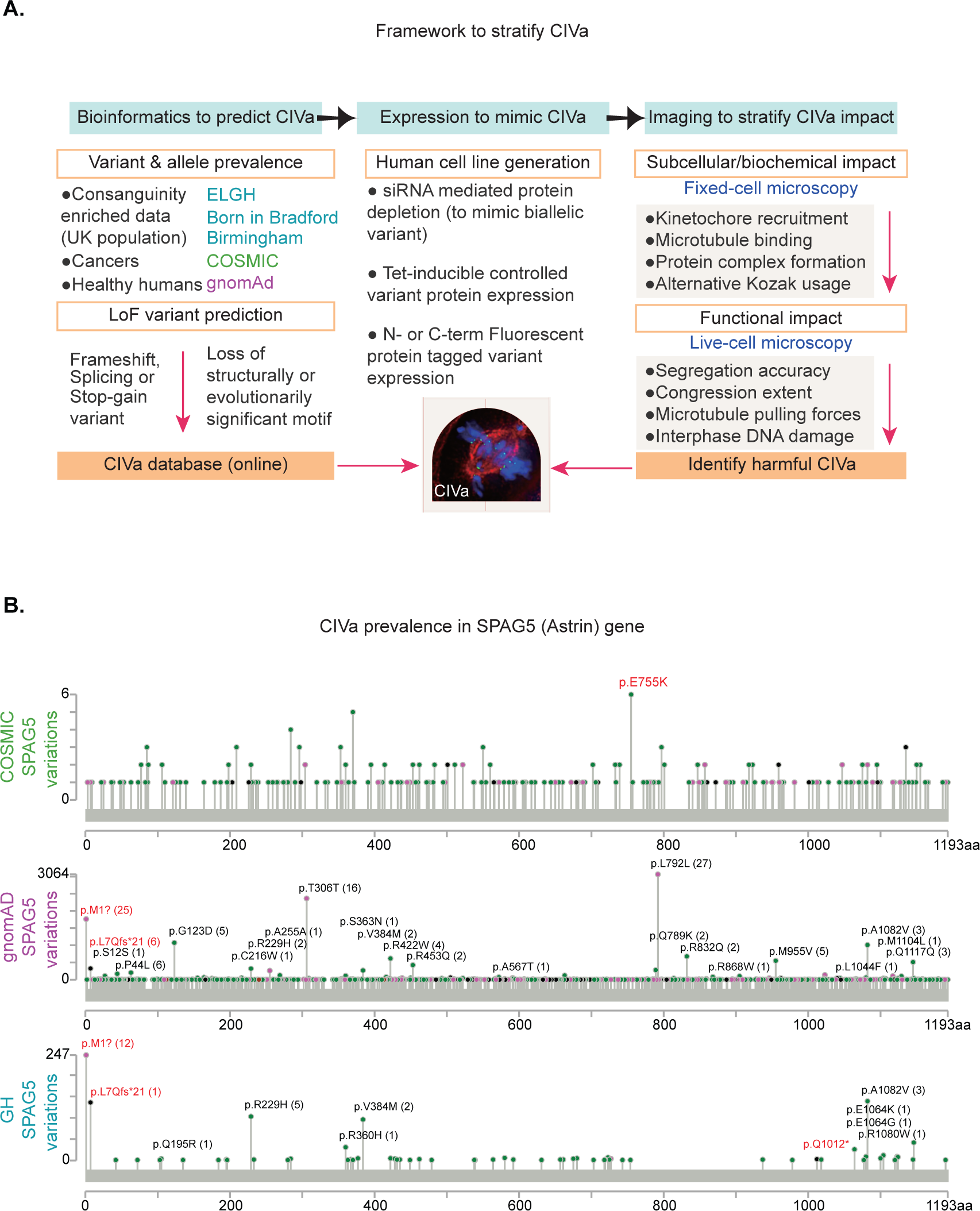
Chromosomal Instability Variant (CIVa) stratification framework. **A**. Cartoon of a three step framework used to predict, mimic and stratify CIVa. CIVa impact stratified through kinetochore localisation and function assessments using microscopy. Bioinformatic predictions on Loss-of-Function variant in kinetochore genes and their allelic prevalence across COSMIC, gnomAD and ELGH databases are collated into the CIVa database. CIVa in kinetochore genes are mimicked in single cells using different protein expression tools and their impact quantitatively assessed to stratify harmful and harmless variants in monoallelic and biallelic forms. **B**. Analysis of CIVa prevalence in SPAG5 gene that encodes Astrin protein using its variation spectrum in COSMIC, gnomAD and ELGH databases. The images show the positions and number of occurrences of different types of variations in the SPAG5 gene, including potential CIVa sites. A black dot indicates a truncating mutation, a green dot indicates a missense mutation and a purple dot indicates other mutation types. The red font indicates interesting variations. Numbers in brackets indicate the number of homozygous occurrences.

To collate a CIVa dataset, we first surveyed for predicted loss-of-function (LoF) variants in chromosome segregation genes in the GH database^11^ of individuals with a high incidence of consanguineal marriages increasing the chances of identifying gene knock-outs directly in humans^12^ (heterozygous and homozygous variants listed in Table 1). Out of the 141 CIVa predictions identified, only 6 are found as homozygous (biallelic) variants, suggesting poor tolerance to CIVa. Unexpectedly, we identified two LoF variants in the SPAG5, p.(Q1012*) and p.(L7Qfs*21), and a highly prevalent LoF variant in SKA3, p.(Q70Kfs*7), proteins crucial for the accurate segregation of chromosomes^13,14,15,16^ (Table 1).

To ask whether CIVa identified in the GH database are specific to the population surveyed, we screened wider population databases: COSMIC (database of cancer patients)^17^, gnomAD (coalition of population-specific and disease-specific databases)^18^ and 1000 Genomes Project (International genome sample resource) (Fig. 1A). The Astrin p.(Q1012*) variant is absent in both COSMIC and gnomAD (Fig. 1B), suggesting it may be specific to the Pakistani-Bangladeshi community surveyed by GH database (Table 1). In contrast, the Astrin p.(L7Qfs*21) variant is present in gnomAD as both heterozygous and homozygous forms (Fig. 1B), indicating a wider spread. Five non-South Asians (European, African-American) with the variant are listed on gnomAD^19^ and the variant is exclusive to South Asians (allele frequency of 2%) in the 1000 Genomes Project Phase 3^20^. We conclude that some of the CIVa identified in the GH database are present in non-south Asian populations.

Using variants in multiple tumour tissues, we assessed the incidence of somatic mutations in Astrin gene by comparing five categories: a) MCPH genes, b) MVA genes c) the Astrin-SKAP complex d) the Astrin-SKAP complex interacting partners and e) TP53 and BRCA1 (two tumour suppressor genes, selected as positive controls). As expected, mutations in the TP53 gene were found in all tissues, including the Gastrointestinal tract, Placenta and Pleura (Fig. 1S1). The percentage of cases with TP53 mutation was above 50% in most of the analysed tumour types, showing a high prevalence of TP53 mutation (Fig. 1S1). In contrast, mutations in the BRCA1 gene, MCPH genes and most of the Astrin-SKAP complex interacting partners, including Ndc80^14,21^, were not ubiquitously present suggesting tissue-specificity (Fig. 1S1). Moreover, the frequency of mutations in the Astrin-SKAP complex and its interacting Ndc80 complex were much lower compared to MCPH or BRCA1 genes, indicating a lower incidence of somatic LoF mutations in Astrin-SKAP genes (Fig. 1S1, see box). The most frequently observed mutation in Astrin p.E755 converted to a K (charge conversion) is specifically found in skin cancers (n=3, Fig. 1B). Thus the percentage of tumour samples with somatic mutations in Astrin-SKAP complex, NDC80 complex and MVA genes is not high, highlighting the uniqueness and potential significance of the two LoF Astrin gene variants, p.(Q1012*) and p.(L7Qfs*21). To allow similar comparisons of genetic and somatic CIVA in 135 chromosome segregation genes, we include a queryable online interface: https://github.com/Draviam-lab/CIVa.

The Astrin p.(Q1012*) variant (Fig. 2A) is predicted to lack a C-terminal tail required for Astrin’s mitotic localisation and function^14^. To determine the subcellular localisation of Astrin p.Q1012*, we expressed YFP-tagged Astrin wild-type (WT) or p.Q1012* in the human epithelial cell line, HeLa (Fig. 2S1 A). Immunostaining show that while Astrin WT localises at the kinetochores (as crescents) and along spindle microtubules, Astrin p.Q1012* does not localise at the kinetochores but is found normally along spindle microtubules (Fig. 2S1 B-C). We compared the severity of Astrin p.Q1012*’s localisation defects against two previously reported Astrin C-terminal mutants (Fig. 2S1 B), either lacking a phosphatase docking motif (RVMF to AAAA, termed 4A mutant) and or lacking the entire C-terminal tail (Δ70 mutant), which promote chromosome missegregation^14^. Quantitative analysis of Astrin crescents showed that the kinetochore enrichment of Astrin p.Q1012* is impaired similarly to the Astrin Δ70 mutant, and more severely than the 4A mutant (Fig. 2S1 C-D), confirming p.Q1012* as a severe loss of kinetochore localisation variant.

**Figure 2.**
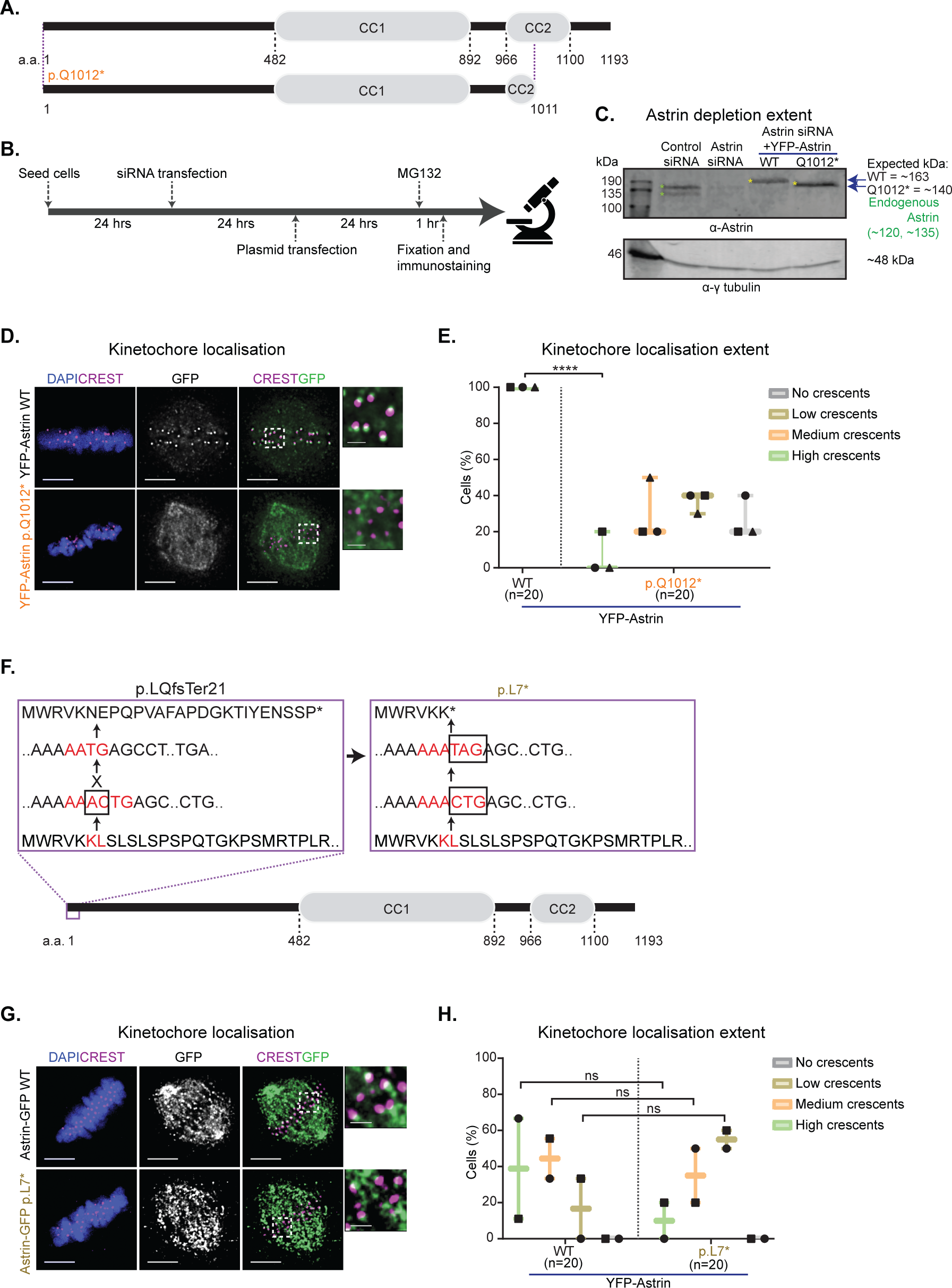
CIVa in Astrin affect kinetochore but not spindle localisation. **A**. Cartoon showing Astrin p.Q1012* variant. **B**. Experimental regimen. **C**. Immunoblot showing Astrin depletion extent in cells treated as in B. **D**. Representative immunofluorescence images of Astrin wild type and p.Q1012* expressing cells treated as in B and probed for GFP and CREST. DNA was stained with DAPI. Scale bars: 5 µm in uncropped images and 1 µm in insets. **E**. Box plot showing Astrin localization at the kinetochores (scored as high, medium, low and no crescents). Symbols represent independent experiments. Two-way ANOVA with Sidak correction was performed for statistical significance. ‘****’ represents p<0.0001. **F**. Cartoon showing Astrin p.L7Qfs*21 variant and Astrin p.7*. **G**. Experimental regimen. **H**. Representative immunofluorescence images of wild type and Astrin p.7* expressing cells treated as in G and probed for GFP and CREST. DNA was stained with DAPI. Scale bars: 5 µm in uncropped images and 1 µm in insets. **I**. Box plot showing Astrin localization at the kinetochores (scored as high, medium, low and no crescents). Symbols represent independent experiments. A chi-square test was performed for statistical significance. ‘****’ represents p<0.0001.

We investigated whether competition with endogenous full-length Astrin prevents the Astrin p.Q1012* variant from localising at the kinetochore. We depleted endogenous Astrin using siRNA and expressed a siRNA resistant form of Astrin p.Q1012* (Fig. 2B-C). Following the depletion of endogenous Astrin, only ∼5% of cells expressing YFP-Astrin p.Q1012* display normal kinetochore-associated crescents (Fig. 2D-E). In conclusion, although Astrin p.Q1012* variant localises normally along microtubules, its kinetochore localisation is severely impaired, both in the presence and absence of endogenous Astrin.

To study the localisation of Astrin variant, p.(E755K), found in skin cancers, we expressed YFP-tagged Astrin E755K, in the presence of endogenous Astrin (Fig. S1 E). Astrin p.E755K localises normally along microtubules and kinetochores (Fig. S1 F-G), indicating that the p.E755K variant does not disrupt Astrin’s role in sensing or maintaining end-on attachments^14,22^.

We mimicked Astrin p.(L7Qfs*21) variant using a C-terminal YFP-tagged Astrin with a stop codon at 7 a.a (Fig. 2F). In the presence of endogenous Astrin, Astrin-YFP wild-type (positive control) and p.L7* mutant localised normally along spindle microtubules (Fig. 2G). As reported previously^14^, fusing GFP at the C-terminus of Astrin wild-type (WT) reduced its localisation at the kinetochores (Fig. 2H).

Importantly, the kinetochore localisation of the Astrin p.L7* mutant was more severely compromised compared to the WT control, but not completely lost from the kinetochore (Fig. 2H), suggesting that a shorter isoform of Astrin WT may be expressed to localise it at the kinetochore.

Unexpectedly, the Astrin LoF variant p.(L7Qfs*21) is found as a homozygous and heterozygous form with a relatively high frequency (Table-1). To test whether the variant allows shorter Astrin, we used bioinformatic tools to predict KOZAK sequences ((gcc)gccRccAUGG), and identified two alternative translation start sites in Astrin p.(L7Qfs*21) at N-454 and N-823. To mimic these start sites, we generated Astrin N-terminal deletion mutants, Δ151 and Δ274, tagged with YFP at their N-termini (Fig. 3A) and expressed them in HeLa cells to assess corresponding protein sizes using immunoblotting. Both Δ151 and Δ274 mutants migrated as expected at 144 kDa and 131 kDa, respectively (Fig. 3B). Importantly, YFP-Astrin Δ151 migrated similarly to Astrin p.L7*-GFP, revealing that Astrin p.(L7Qfs*21) can promote the expression of an N-terminally truncated protein starting from 152 a.a. (N-454) of Astrin.

**Figure 3.**
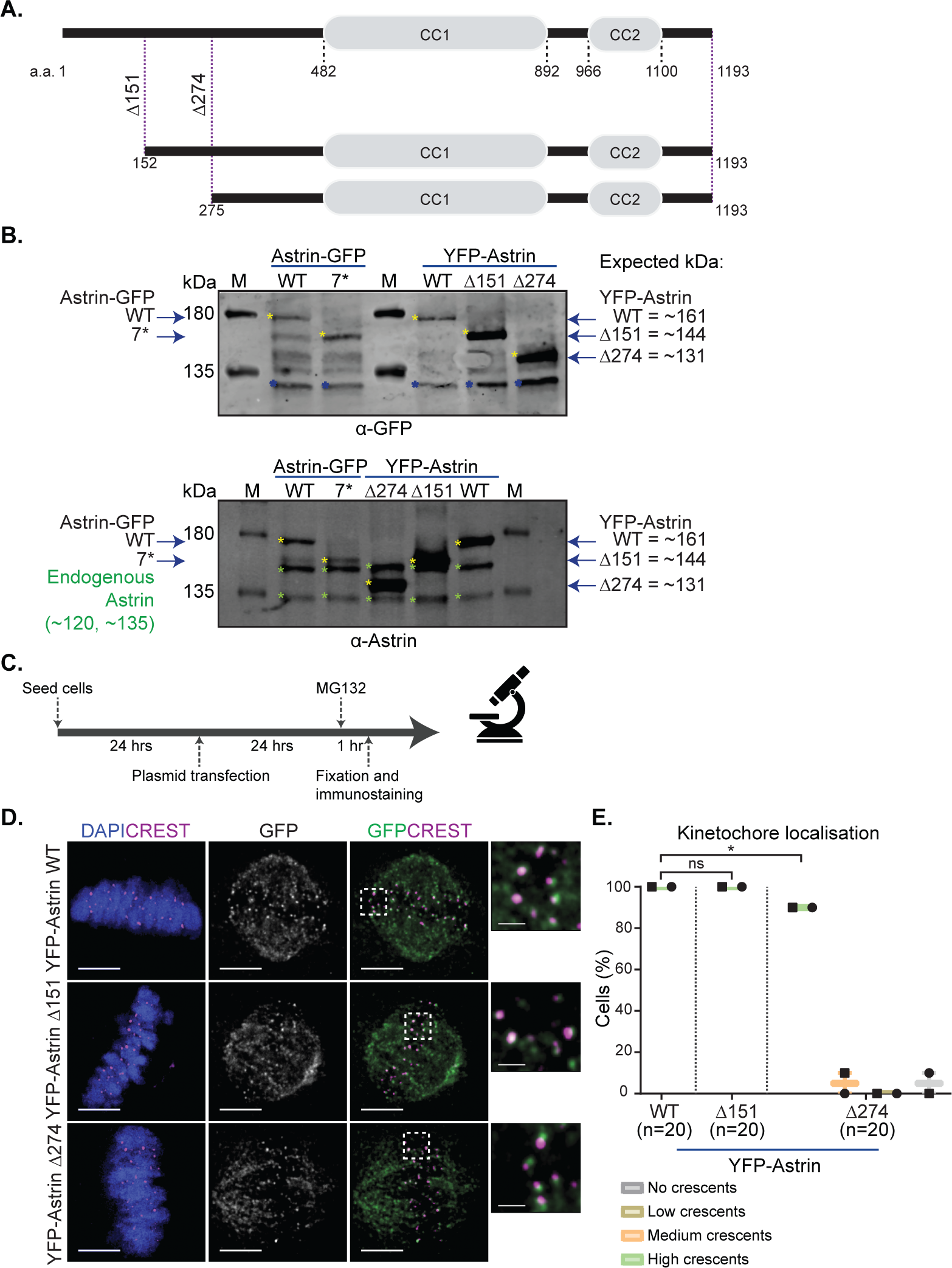
Shorter protein provides resilience against N-terminal CIVa and allows kinetochore localisation. **A**. Cartoon showing N-terminal Astrin mutants. **B**. Immunoblots of HeLa cell lysates expressing Astrin-GFP (wild type and p.7*) and YFP-Astrin (wild type, Δ151 and Δ274) and probed for GFP and Astrin. Yellow, green and blue asterisks refer to GFP fusion protein specific, endogenous Astrin specific and non-specific bands, respectively. **C**. Experimental regimen. **D**. Representative immunofluorescence images of Astrin wild type, Δ151 and Δ274 cells treated as in C and probed for GFP and CREST. DNA was stained with DAPI. Scale bars: 5 µm in uncropped images and 1 µm in insets. **E**. Box plot showing Astrin localization at the kinetochores (scored as high, medium, low and no crescents). Symbols represent independent experiments. Two-way ANOVA with Sidak correction was performed for statistical significance. ‘*’ and ns represent p<0.05 and ‘not significant’ respectively.

As C-terminal tags interfere with Astrin’s kinetochore localisation (Fig. 2G), we tested the subcellular localisation of N-terminal YFP tagged Astrin Δ151 and Δ274 mutants. Immunostaining showed that in the presence of endogenous Astrin, both Astrin Δ151 and Δ274 localise normally along spindle microtubules (Fig. 3D). While Δ151 mutant localises normally at the kinetochore, Δ274 mutant exhibits reduced kinetochore localisation (Fig. 3 C-E). However, in the absence of endogenous Astrin, both Δ274 and Δ151 mutants localised normally at the kinetochore (Fig. 3S1 B-C). Thus endogenous full length Astrin can outcompete the recruitment of Astrin Δ274 deletion mutant at the kinetochore, whereas in the absence of full length protein both Astrin Δ151 and Δ274 mutants localise normally at kinetochores.

Astrin forms a 4-subunit complex^21^ with SKAP as one of the partner^13^, and SKAP requires Astrin for its localisation at kinetochores^13^. To assess if the Δ274 mutant can normally recruit the Astrin-SKAP complex to kinetochores, we depleted endogenous Astrin and analysed SKAP localisation. Following Astrin depletion, the Δ274 mutant localises normally at kinetochores (Fig. 3S1), which serves as a good internal control for endogenous Astrin depletion status. Immunostaining studies of metaphase arrested cells depleted of endogenous Astrin and expressing Δ274 mutant, showed normal kinetochore localisation of SKAP (Fig.3S1 B-C). Thus shorter forms of Astrin (Δ274) can localise at the kinetochore, and recruit SKAP normally, consistent with previous studies^13,23^.

Both Astrin Δ274 and Astrin Δ151 are normally excluded from the interphase nucleus and present at microtubule-ends, suggesting normal interphase localisation and function (Fig. 3S1 D). We conclude that the Astrin p.(L7Qfs*21) variant is likely to express a shorter Astrin that lacks the first 151 a.a of Astrin, but can localise normally at kinetochores, explaining the wide prevalence of this variant and revealing a resilience mechanism that copes with N-terminal LoF variants.

Another N-terminal LoF variant is frequently observed in SKA3, p.(Q70Kfs*7) variant (1906 heterozygous and 2 homozygous)^11^, in a region essential for the SKA complex formation^24^. This variant has been reported in multiple cancers including Haematopoietic and lymphoid, Thyroid, Pancreas, Oesophagus and Lung^17^ (Table 1). To study the variant’s localisation, we generated GFP-SKA3 p.Q70Kfs*7 and coexpressed with a CENPB-dsRed (a centromere marker). Unlike GFP-SKA3 WT, the GFP-SKA3 p.Q70Kfs*7 fails to localise at kinetochores during mitosis (Fig. 3S2 A). Similarly, unlike C-terminal CFP-tagged SKA3 WT, SKA3 p.Q70Kfs*7-CFP fails to localise at kinetochores (Fig. 3S2 B). Immunostaining studies using antibodies against 156-177 a.a of SKA3 showed that in cells expressing GFP-SKA3 p.Q70Kfs*7, endogenous SKA3 localises normally at kinetochores (Fig. 3S3 A and B). In mitotic cells expressing either SKA3 p.Q70Kfs*7 fusion proteins, we observed normal chromosome congression, and hence we probed if mature kinetochore attachments had formed using the end-on attachment marker, Astrin-SKAP complex^5^. Live-cell microscopy of cells coexpressing mKate2-Astrin and SKA3 showed normal Astrin localisation in both N-terminally and C-terminally tagged SKA3-WT and Q70Kfs*7 expressing cells (Fig. 3S2 C and D). In summary, we find that the SKA3 Q70Kfs*7 does not interfere with metaphase chromosome congression or end-on attachment status, suggesting normal chromosome-microtubule attachments in cells expressing SKA3 p.(Q70Kfs*7) variant, revealing a different type of resilient mechanism to cope with an N-terminal LoF variant.

The Astrin p.(Q1012*) variant is unique with impaired kinetochore localisation (Fig. 2 and 3). As the Astrin-SKAP complex is observed as a dimer^21^, we hypothesised that Astrin pQ1012* expression may disrupt the kinetochore localisation of the endogenous Astrin-SKAP complex. To test this hypothesis, we expressed Astrin wild-type and p.Q1012* in HeLa cells and analysed the kinetochore localisation of endogenous Astrin and SKAP using immunostaining studies (Fig. 4S1 A). Astrin localises as a crescent at only 68% of Astrin p.Q1012* expressing cells compared to 96% of cells expressing Astrin wild type (Fig. 4S1 B-C). Similarly, endogenous SKAP localised at only 86% of Astrin p.Q1012* expressing cells compared to 97% of wild type expressing cells (Fig. 4S1 D-E). Thus the expression of Astrin p.Q1012* can disrupt the kinetochore localisation of endogenous full-length Astrin-SKAP complex in a dominant-negative manner.

Astrin’s C-terminal tail which is lost in Astrin p.Q1012* serves two important roles: it delivers PP1 phosphatase to the kinetochore which stabilises end-on kinetochore-microtubule (MT) attachments and it enables microtubule-mediated pulling which ensures maximum enrichment of Astrin-SKAP at kinetochores^14,23,25^. So, we investigated the extent to which microtubule-mediated pulling is reduced and Astrin p.Q1012* localisation is reduced following Astrin p.Q1012* expression by measuring inter-centromeric distances in cells co-expressing CENPB-DsRed, a centromere marker, along with YFP-tagged Astrin WT or pQ1012* mutant (Fig. 4 S2 A-B). The range of inter-centromere distances was reduced in cells expressing YFP-Astrin p.Q1012* compared to YFP-Astrin WT (Fig.4 S2 B-D). This difference in inter-centromere distances was more striking following the normalisation of inter-centromere distances of each pair to its unstretched state (marked as T_0_). Tracking the fate of sister kinetochores for 5 minutes showed that although centromeric stretching can be observed in Astrin p.Q1012* expressing cells, the maximum inter-centromeric distances are reduced compared to Astrin WT expressing cells (Fig. 4 S2 E-F). We conclude that stable microtubule-mediated pulling of kinetochores is reduced in cells expressing the Astrin pQ1012* variant, despite the presence of endogenous full-length Astrin.

Quantifying the extent of loss of kinetochore-bound Astrin p.Q1012* relative to WT in metaphase kinetochores through time required automated analysis. We developed a computational workflow for image analysis using segmentation and particle tracking tools (details in methods). Using the centromeric marker (CENPB-DsRed intensities) as a mask, we measured the intensities of the Astrin p.Q1012* and WT protein in dynamically stretching sister kinetochores (Fig. 4 S3A). While Astrin WT signals at kinetochore were on average 1.1-1.3 fold higher than signal intensities in the cytoplasm, the variant was only 0.9 fold higher relative to cytoplasmic signal intensities, indicating an active reduction of Astrin p.Q1012* at the kinetochore (Fig. 4 S3 B). Tracking these changes through time showed a steady reduction in p.Q1012* associated kinetochore intensities (Fig. 4 S3C). Thus, the severe reduction of p.Q1012* at kinetochores correlates well with the reduced microtubule-mediated pulling of sister kinetochores in Astrin p.Q1012* expressing cells.

To investigate the fate of mitotic cells expressing the p.Q1012* variant, we generated a tetracycline-inducible HeLa FRT/TO YFP-Astrin Q1012* cell line and acquired images every 6 minutes for 10 hrs in the presence of SiR-DNA (a DNA tracker) following a brief exposure to Tetracycline (Fig. 4A). Quantitative analysis of time-lapse movies showed that only 70% of Astrin p.Q1012* expressing cells completed mitosis compared to 95% of Astrin WT expressing cells (Fig.4S4 A). Astrin p.Q1012* expressing cells that completed mitosis displayed a delayed anaphase onset (AO).

**Figure 4.**
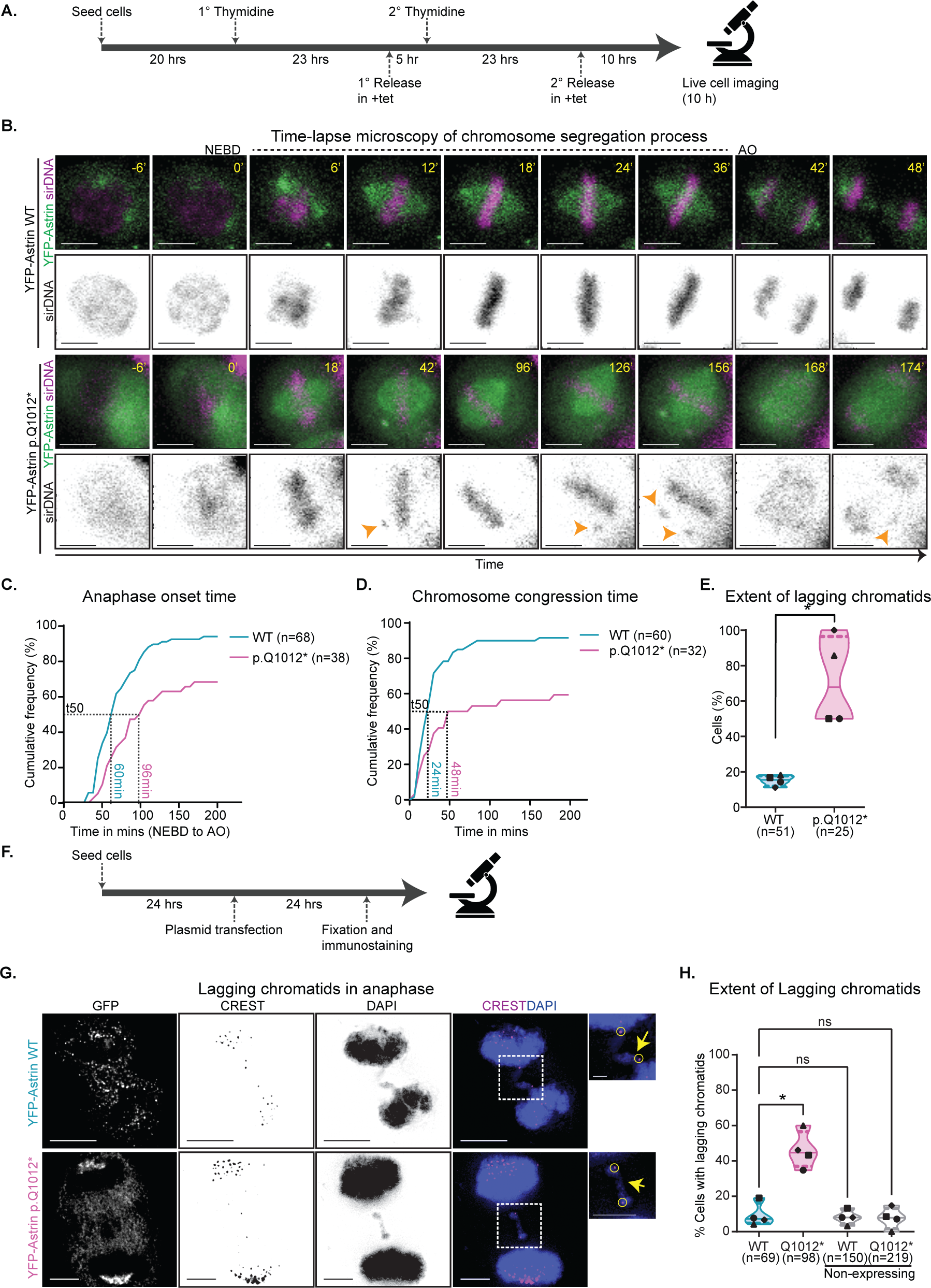
Astrin p.Q1012* variant prolongs mitosis and promotes lagging chromosomes. **A**. Experimental regimen. **B**. Representative time-lapse images of Astrin wild type and p.Q1012* cells treated as in A. Arrows represent lagging chromosomes. NEBD is nuclear envelope breakdown and AO is anaphase onset. Scale bars: 15 µm. **C**. Cumulative frequency graph showing the time taken from NEBD to AO. T50 indicates the time taken by 50% of cells to complete mitosis. **D**. Cumulative frequency graph showing the time taken from NEBD to the formation of the metaphase plate. T50 indicates the time taken by 50% of cells to congress chromosomes. **E**. Violin plot showing the percentage of cells with lagging chromosomes at anaphase. The solid line represents the median and dotted lines represent the quartiles. Each dot represents an independent set. Paired t-test was performed to find statistical significance. ‘*’ represents p<0.05. (C-E) Data represents four independent sets. **F**. Experimental regimen **G**. Representative immunofluorescence images of Astrin wild type and p.Q1012* expressing anaphase cells with lagging chromosomes. Cells were treated as in F and probed for GFP and CREST. DNA was stained with DAPI. Scale bars: 5 µm in uncropped images and 1 µm in insets. **H**. Violin plot showing the percentage of anaphase cells with lagging chromosomes. The solid line represents the median and dotted lines represent quartiles. Each dot represents an independent experiment. One-way ANOVA with DUNNET correction was performed to find statistical significance. ‘*’ and ns represent p<0.05 and ‘not significant’, respectively.

To accurately quantify AO times, Nuclear Envelope Break Down (NEBD) was ascertained by the loss of the nucleo-cytoplasmic boundary of cytoplasmic YFP-Astrin signals. The time from NEBD to AO was 1.5 fold longer in YFP-Astrin p.Q1012* expressing cells compared to YFP-Astrin WT expressing cells (96 mins for p.Q1012* *versus* 60 mins for WT), showing significant delay in anaphase onset (Fig. 4B-C). Both the delay in anaphase onset and increased incidence of mitotic failure indicate that the expression of Astrin p.Q1012* variant promotes a chromosome segregation defect.

To investigate the underlying cause for the prolonged mitosis in Astrin p.Q1012* expressing cells, we analysed chromosome congression in the time-lapse movies. We found that only 56% of Astrin p.Q1012* expressing cells successfully congressed their chromosomes compared to 90% of Astrin WT expressing cells (Fig. 4A, D). Additionally, cells expressing Astrin p.Q1012* are two-fold slower in chromosome congression compared to those expressing Astrin WT; (t50 of NEBD to metaphase: 48 mins for Q1012* *versus* 24 mins for WT; Fig. 4A, D). Importantly, once congressed, only 30% of Astrin p.Q1012* expressing cells maintain chromosome congression compared to 76% of Astrin WT expressing cells (Fig. 4S4 B). Thus, both the congression of chromosomes onto a metaphase plate and the maintenance of a congressed plate are disrupted in YFP-Astrin Q1012* expressing cells, confirming errors in stable chromosome-microtubule attachment.

To investigate whether the prolonged mitosis, reduced microtubule pulling and chromosome congression defects induced by Astrin p.Q1012* have an impact on chromosome segregation accuracy, we scored the presence of lagging chromosomes in anaphase cells. Time-lapse movies showed that 70% of all Astrin p.Q1012* expressing cells presented lagging chromosomes during anaphase compared to 15% of Astrin WT expressing cells (Fig. 4A, E). Moreover, immunostaining studies showed that ∼46% of Astrin p.Q1012* expressing anaphase cells display lagging chromatids compared to ∼9% in Astrin WT expressing cells (Fig. 4 F-H). Thus, Astrin p.Q1012* variant significantly increases the incidence of missegregating chromosomes and lagging chromatids during anaphase. We conclude that the expression of the naturally occurring variant Astrin p.(Q1012*) despite the presence of Astrin full length protein (as in heterozygous/monoallelic form) is likely to promote DNA damage and chromosomal instability in humans.

## DISCUSSION

We present the first comprehensive survey of <u>C</u>hromosomal Instability aiding <u>V</u>ariants (CIVa) in chromosome segregation genes by assessing LoF variant frequency across multi-ethnic populations. We develop single-cell studies to systematically analyse the significance of CIVa predictions in two essential chromosome segregation genes and show the existence of resilience mechanisms to cope with some but not all harmful variants. Astrin p.Q1012* variant impairs the localisation of endogenous full-length Astrin in a dominant negative manner, leading to chromosome segregation defects. In contrast, Astrin p.L7Qfs*21 results in the expression of a shorter fragment of Astrin lacking the N-terminal region but localises normally at microtubule-ends and kinetochores. Our findings may explain the relatively higher prevalence of Astrin p.(L7Qfs*21) but not p.(Q1012*) variant in the general population. We show that another highly prevalent monoallelic variant of SKA3 p.(Q70Kfs*7) does not localise at kinetochores or interfere with endogenous SKA3 localisation. Thus, systematic assessment of the impact of CIVa will allow the interpretation of the impact of homozygous or compound heterozygous LoF genetic variations predicted to promote CIN.

Previous evaluation of Astrin variants (p.[(G1064E*3)];[(K409Pfs*19)]) known to present clinical features (microcephaly) were limited to compromised centrosomal localisation of Astrin^6^. We present the first assessment of Astrin variants’ loss of kinetochore localisation and its impact on chromosome segregation. Of the two LoF Astrin variants, we identified, Astrin p.(Q1012*) (heterozygous=3; GH database) might be exclusive to the Pakistani-Bangladeshi community in East London^11,17,18^. In contrast, Astrin p.(L7Qfs*21) variant is found across multiple ancestries (heterozygous=134, homozygous=1; GH database), (heterozygous=13, homozygous=0; GenomeAsia 100K) and (heterozygous=320; homozygous=6; gnomAD database)^11,17,18^. Moreover, a homozygous Astrin start loss variant is listed on the GH (heterozygous=223; homozygous=12), gnomAD (heterozygous=1761; homozygous=25), GenomeAsia 100K (heterozygous=21; homozygous=0) and UK10K databases^11,18,20^, i.e., present in all ethnicities with a higher incidence among Europeans^18^. Astrin start-loss and p.(L7Qfs*21) variants are the only homozygous LoF Astrin variants found in the gnomAD database^18^. Heterozygous Astrin start loss and p.(L7Qfs*21) variants are also listed on the GenomeAsia100K and TOPmed databases^26,27^. We speculate that the two homozygous variations in the N-terminus of Astrin may have originated in two continents.

The SKA3 variant p.(Q70Kfs*7) is predicted to disrupt the multimerisation domain within the 3 subunits (SKA1-SKA2-SKA3) of the SKA complex^24^. However, we find that endogenous SKA3 localisation is unperturbed in cells expressing SKA3 p.Q70Kfs*7, suggesting a reason for the variant’s prevalence in heterozygous form. In contrast, Astrin p.Q1012* is absent at end-on kinetochores impairing Astrin’s vital kinetochore-microtubule bridging function and promoting chromosome missegregation. Thus, a framework to combine human genetic variant prevalence and function impact in dividing cells can help stratify CIVa relevant to a variety of pathologies.

## METHODS

### Genome Databases and Acces

To curate the CIVa dataset, we have used multiple public databases. 1000 Genomes project Phase 3, UK10K database, the GH database (sample size=8,921), The Catalogue of Somatic Mutations in Cancer (COSMIC) database (sample size>37,000) and gnomAD (sample size=141,456) were all assessed on: Jan 2022. Table-1 was built using the GH LoF variants list from 2018 and the lollipop graph was generated using GH all variants data 2020 (both accessed on 29.01.2020). Lollipop graphs were also generated using data from gnomAD v.2.1.1 and COSMIC databases (assessed on 29.01.2020). Cancer mutational spectra heatmap was generated from data using the COSMIC database (assessed on 29.01.2020). Lollipop graphs were generated using MutationMapper (cBioPortal for Cancer Genomics::MutationMapper, assessed on 29.01.2020). The list of Chromosome segregation genes in Table1 was identified using the Gene Ontology term ‘Kinetochore’ (GO:0000776). For Table-1, gnomAD (v2.1.1) and COSMIC databases (v5) were accessed in Jan 2022.

### Cell Culture, Plasmids and Transfection

HeLa cells (ATCC) were cultured in Dulbecco’s Modified Eagle’s Media (DMEM) supplemented with 10% FCS and antibiotics (Penicillin and Streptomycin). HeLa FRT/TO YFP-Astrin cell lines were cultured in tetracycline free DMEM supplemented with 10% FCS and antibiotics. YFP-Astrin p.Q1012*, YFP-Astrin p.E755K and Astrin-GFP p.L7* expression plasmids were generated by site-directed point mutagenesis. YFP-Astrin Δ151 and Δ274 expression plasmids were generated by amplifying regions 152-1193 a.a. and 275-1193 a.a. respectively and subcloning into a YFP expression plasmid. YFP-Astrin WT and Astrin-GFP WT expression plasmids were previously described^14^. pcDNA5 FRT/TO YFP-Astrin p.Q1012* expression plasmid was generated by subcloning YFP-Astrin p.Q1012* into pcDNA5 FRT/TO plasmid. A single Cytosine nucleotide (position 208) deletion mutant of SKA3 fused to GFP or CFP to generate GFP-SKA3 p.Q70Kfs*7 or SKA3 p.Q70Kfs*7-CFP expression plasmids, respectively. Plasmid sequences were confirmed by DNA sequencing. siRNA transfection was performed using Oligofectamine according to the manufacturer’s instructions. To target Astrin mRNA, Astrin 52 oligo (UCCCGACAACUCACAGAGAAAUU) was used. Negative control siRNA (12,935–300) was from Invitrogen. Plasmid transfection was performed using TurboFect (Fisher; R0531) or DharmaFECT duo (Dharmacon; T-2010) according to the manufacturer’s instructions. In addition to the standard protocol, after 4 h of incubation, the transfection medium was removed and a fresh selected pre-warmed DMEM was added to each well. In co-transfection studies, eukaryotic expression vectors encoding Astrin and CENPB were used in a 3:1 ratio. HeLa FRT/TO YFP-Astrin p.Q1012* cell line was generated by transfection of HeLa Flp-In cells with pCDNA5-FRT/TO-YFP-Astrin p.Q1012* expression plasmid followed by a brief Hygromycin selection and sorting for YFP positive cells using FACS. HeLa FRT/TO YFP-Astrin wild type cell line was previously described^14^. Induction of exogenous YFP-Astrin was performed by exposing the cells to DMEM medium supplemented with Tetracycline. For localization and inter-centromeric distance studies, cells were treated with 10□µM MG132 (TOCRIS; 1748) for one hour. For the mitotic progression study, cells were synchronised using 2.5 mM Thymidine (ACROS organics).

### Immunostaining studie

Cells were cultured on ø13 mm round coverslips (VWR; 631-0150) and fixed with ice-cold methanol for one minute. Following fixation, two quick washes with a PBST wash buffer (1X PBS + 0.1% Tween 20) were performed, followed by two washes of 5 minutes each. Coverslips were incubated with (1X PBS + 0.1% Tween 20 + 1% BSA) for 20 minutes, before staining with primary antibodies overnight at 4°C followed by two washes before incubation with secondary antibodies for 30 minutes at room temperature. Finally, coverslips were washed twice with PBST except before mounting onto glass slides when coverslips were quickly rinsed in distilled water.

Cells were stained with antibodies against GFP (Roche; 1181446001; 1:1000), GFP (Abcam; ab290; 1:1000), SKAP (Atlas; HPA042027; 1:800), Astrin (Proteintech; 14726–1-AP; 1:1000), SKA3 (Santa Cruz; H-9; 1:500) and CREST antisera (Europa; FZ90C-CS1058; 1:2000). DAPI (Sigma) was used to stain DNA. All antibody dilutions were prepared using the blocking buffer. Images of immunostained cells were acquired using 100X/NA1.4 UPlanSApo oil immersion objective on a DeltaVision Core microscope equipped with CoolSnap HQ Camera (Photometrics). Deconvolution of fixed-cell images was performed using SoftWorx™.

### Live-cell Imagin

For live-cell imaging studies, cells were seeded onto 4-well cover glass chambered dishes (Lab-Tek; 1064716) and transferred to Leibovitz’s L15 medium (Invitrogen; 11415064) for imaging. For low-resolution live-cell imaging, HeLa FRT/TO YFP-Astrin cells were synchronised using a double thymidine block. 100 nM sirDNA (Tebu-bio; SC007) was added 10 hours before image acquisition to stain for DNA. 3 *Z*-planes, 0.6□μm apart, were acquired using a 40X/0.95 UPlanSApo air objective on an Applied Precision DeltaVision Core microscope equipped with a Cascade2 camera under EM mode. Imaging was performed at 37°C using a full-stage incubation chamber set up to allow normal mitosis progression and microtubule dynamics. For high-resolution live-cell imaging, cells were transfected with plasmid vectors 24□hours before an hour-long 10□µM MG132 treatment to arrest mitotic cells in metaphase. 3 *Z*-planes, 0.6□μm apart, were acquired using a 100X/1.40 UPlanSApo oil immersion objective on an Applied Precision DeltaVision Core microscope equipped with a Cascade2 camera under EM mode. For live-cell CFP imaging, Applied Precision DeltaVision Elite microscope equipped with an EDGE sCMOS_5.5 camera with a 60X oil-immersion objective was used. Imaging was performed at 37°C using a full-stage incubation chamber set up to allow normal mitosis progression and microtubule dynamics. SoftWorx™ distance measurement tool was used to find inter-centromeric distances. Additional analysis was conducted on Microsoft Excel and graphs plotted using GraphPad Prism 9™.

### Immunoblotting studie

Immunoblotting was performed on proteins separated on 8% or 12% SDS-PAGE gels by transferring them overnight onto Nitrocellulose membrane. Membranes were incubated in primary antibodies against Astrin (Proteintech; 14726–1-AP; 1:3000), γ-Tubulin (Sigma-Aldrich; T6793; 1:800), GFP (Abcam; ab290; 1:1000) and SKAP (Atlas; HPA042027; 1:1000), and probed using secondary antibodies labelled with infrared fluorescent dyes, which were imaged using an Odyssey (LiCOR) imager.

### Kinetochore Particle Tracke

The image processing tool to measure kinetochore signal intensities was developed in Python 3, using python’s image processing libraries scikit-image in Anaconda Environment and Jupyter Notebook. Data analysis was done in RStudio with the package ggplot. Fig.ure panels were generated using matplotlib, ggplot and jupyter-notebook. Initial image pre-processing was done in ImageJ. To measure the kinetochore intensities in 3D images of time-lapse movies, the CENPB-dsRed signal was first detected to identify the location of kinetochores by applying an edge detector filter and a suitable threshold. Small particles were removed, and the holes were filled by performing morphological operations. Next, by burning the CENPB signal mask on the YFP-Astrin channel image, we extracted the mean particle intensities of YFP-Astrin. The cytoplasmic intensity was measured by creating a binary mask to segment the cell from the background and identifying a ring-shaped region as a proxy for the cytoplasm. The source code is available for download at Github: https://github.com/Draviam-lab/Kinetochore-Particle-Tracker

### CIVa database

CIVa database was developed in GitHub Pages using JavaScript, HTML and CSS. This database can be queried on the Gene Symbol or the Uniprot ID. The Gitpage is available at: https://draviam-lab.github.io/CIVa/. The source code and downloadable CIVa dataset is available at Github https://github.com/Draviam-lab/CIVa.

## Acknowledgement

We acknowledge funding support from BBSRC (R01003X/1 and T017716/1 to VMD), QMUL (SBC8DRA2 and SBC9DRA2 to VMD), Chinese Scholarship Council (CSC file no. 201906820034 to XS), CONACYT Scholarship (CVU no.1042679 to JCMG) and CRUK (C28598/A9787 to VMD). We acknowledge David Dang for support with particle tracker studies; Asad Islam for supporting AI’s data curation and file conversion efforts; Sam Court and Petra Ungerer for infrastructure maintenance support; Christoforos Efstathiou and Sophie Adams for detailed comments on the manuscript; and other Draviam group members for discussions on data acquisition and analysis. We thank David Van Heel for his contributions to check the genomic variant sequences reported for Astrin and SKA3 in the ELGH database.

## Author Contributions

AI performed the experiments, analysed the data and generated the panels for all main Figures except for Figure 1A and Supplementary Figures 1S, 2S1, 3S1 A-C, 4S1, 4S2 and 4S4; JCM-G performed the experiments, analysed the data and generated the panels for Figures 3S1 D, 3S2, 3S3 and 3S4; TG performed data analysis and generated the Figure panels for Fig. 4 S3. XS and VMD prepared Figure 1A. VMD conceptualised the study, planned the experiments. The manuscript text was drafted by VMD and AI, and edited by VMD. TG developed the code for the kinetochore particle tracker and CIVa online database. AI, JCMG and XS contributed to data curation for the CIVa database.

## FIGURE LEGEND

**Figure 1S. Cancer mutational spectra**. Heat map of incidence of variations in selected genes in different tumour types in the COSMIC database. Box highlights genes with a low percentage of tumour samples showing somatic mutations.

**Figure 2S1. Astrin variants p.Q1012* and p.E755K localise normally at the spindle but not at the kinetochores in the presence of endogenous Astrin. A**. Experimental regimen. **B**. Cells were transfected with YFP-tagged Astrin (WT) or mutants and fixed for immunostaining following ∼1 hour of MG132 treatment. Cartoon showing Astrin p.Q1012*, 4A and Δ70. **C**. Representative immunofluorescence images of Astrin wild type and p.Q1012*, 4A and Δ70 expressing cells treated as in A and probed for GFP and CREST. DNA was stained with DAPI. Scale bars: 5 µm in uncropped images and 1 µm in insets. **D**. Box plot showing Astrin localization at the kinetochores (scored as high, medium, low and no crescents). Symbols represent independent experiments. Two-way ANOVA with Sidak correction was performed for statistical significance. “****” represents p<0.0001. **E**. Cartoon showing Astrin p.E755K variant. **F**. Representative immunofluorescence images of Astrin wild type and p.E755K expressing cells treated as in A and probed for GFP and CREST. DNA was stained with DAPI. Scale bars: 5 µm in uncropped images and 1 µm in insets. **G**. Box plot showing Astrin localization at the kinetochores (scored as high, medium, low and no crescents). Symbols represent independent experiments.

**Figure 3 S1. Astrin** Δ**274 localises normally at the spindle and kinetochores when endogenous Astrin is depleted. A**. Experimental regimen **B**. Representative immunofluorescence images of YFP tagged Astrin wild type and Δ274 expressing cells treated as in A and probed using antibodies against GFP and SKAP and CREST antisera. DNA was stained with DAPI. Scale bars: 5 µm in uncropped images and 1 µm in insets. **C**. Box plot showing Astrin localization at the kinetochores (scored as high, medium, low and no crescents). Symbols represent independent experiments. **D**. Representative immunofluorescence images of YFP tagged Astrin wild type, Δ151 or Δ274 expressing cells probed using antibodies against GFP and the growing microtubule-end marker, EB1. DNA was stained with DAPI. Scale bars: 5 µm in uncropped images and 5 µm in insets.

**Figure 3 S2. SKA3 p.Q70Kfs*7 does not localise at kinetochores or disrupt end-on attachments. A**. Representative deconvolved images of live mitotic cells coexpressing CENPB-dsRed (centromere marker) and N-terminal GFP tagged SKA3 WT or p.Q70Kfs*7 as indicated (n = 14 WT and 14 p.Q70Kfs*7 mitotic cells). **B**. Representative deconvolved images of live mitotic cells coexpressing CENPB-dsRed (centromere marker) and C-terminal CFP tagged SKA3 WT or p.Q70Kfs*7 as indicated (n = 13 WT and 14 p.Q70Kfs*7 mitotic cells). **C**. Representative deconvolved images of live mitotic cells coexpressing mKate2-Astrin (end-on attachment marker) and N-terminal GFP tagged SKA3 WT or p.Q70* as indicated (n = 16 WT and 19 p.Q70Kfs*7 mitotic cells). **D**. Representative deconvolved images of live mitotic cells coexpressing mKate2-Astrin (end-on attachment marker) and C-terminal CFP tagged SKA3 WT or p.Q70Kfs*7 as indicated (n= 15 WT and 12 p.Q70Kfs*7 mitotic cells). Samples collated from at least 3 independent experimental repeats. Scale bars: 5 µm in uncropped images and 1 µm in insets.

**Figure 4 S1. Endogenous Astrin and SKAP localisation is disrupted in Astrin p.Q1012* expressing cells. A**. Experimental regimen. **B**. Representative immunofluorescence images of Astrin wild type and p.Q1012* expressing cells treated as in A and probed for GFP, Astrin and CREST. DNA was stained with DAPI. Scale bars: 5 µm in uncropped images and 1 µm in insets. **C**. Violin plot showing the percentage of kinetochores where Astrin localised as a crescent. Forty kinetochores were counted per cell. **D**. Representative immunofluorescence images of wild type and p.Q1012* expressing cells treated as in A and probed for GFP, SKAP and CREST. DNA was stained with DAPI. Scale bars: 5 µm in uncropped images and 1 µm in insets. **E**. Violin plot showing the percentage of kinetochores where SKAP localised as a crescent. Forty kinetochores were counted per cell.Dots represent independent cells, the solid line represents the median, dotted lines represent the quartiles and colours represent independent sets. Mann-Whitney U test was performed for statistical significance. ‘****’ and ‘**’ represent p<0.0001 and p<0.01, respectively.

**Figure 4 S2. Inter-kinetochore stretching is reduced in cells expressing Astrin p.Q1012*. A**. Experimental regimen. **B**. Representative time-lapse images of Astrin WT and p.Q1012* expressing cells treated as in A. Scale bars: 5 µm in uncropped images and 1 µm in insets. **C**. Cartoon showing inter-centromeric distance as a measurement of inter-kinetochore (KT) stretching. **D**. Scatter plot showing inter-KT distances in cells treated as in A measured for five pairs of KTs per cell over 10 minutes (1 frame/min). Colours represent different sets. ‘n’ is the number of cells. Error bars show mean with SD. Mann-Whitney U test was performed for statistical significance. ‘**’ represents p<0.001. **E-F**. Change in inter-KT distances (normalised to least inter-KT distance) over time in cells treated as in A. ‘0’ is the time point of least inter-KT distance. The shaded area represents a 95% confidence interval. ‘nKT’ is the number of kinetochores. (D-F) Data represent three independent experiments.

**Figure 4 S3. Kinetochore particle intensities show a steady reduction in Astrin p.Q1012* variant levels compared to WT. A**. Automation routine for measuring YFP Astrin intensities at kinetochore particles. A particle mask was developed using CENPB-ds-Red (a centromere marker) image channel and then burned on the YFP-Astrin image channel. The detected kinetochore particles are then labelled and the mean intensity of Astrin at the particle site is calculated. The particles are coloured by their mean intensity values (Dark blue to light blue: high to low intensity). Scale bar: 5 μm **B**. Violin Plot showing the distribution of Astrin intensities at kinetochore (KT) particles in WT and p.Q1012* expressing cells. The box plot shows the mean kinetochore intensity is less in p.Q1012* than the WT. Mann-Whitney U test was performed for statistical significance. ‘****’ represents p< 0.0001. n refers to the number of cells. **C**. Time-lapse analysis of Astrin intensities. Jitter is added to avoid overplotting. Each jitter point is the mean of the intensity ratio at that time point per cell for each condition.

**Figure 4 S4. A proportion of stably expressing Astrin p.Q1012* cells fail to complete mitosis and cannot maintain chromosome congression. A**. Violin plot showing the percentage of Astrin wild type and Astrin p.Q1012* expressing cells which successfully exited mitosis. Each dot represents an independent experiment. **B**. Violin plot showing the percentage of Astrin wild type and Astrin p.Q1012* expressing cells that successfully maintained chromosome congression. Each dot represents an independent experiment. The solid and dotted lines represent the median and quartiles, respectively. ‘*’ and ‘**’ represents p<0.05 and p<0.01, respectively.

